# Stability-based sorting: The forgotten process behind (not only) biological evolution

**DOI:** 10.1101/149963

**Authors:** Jaroslav Flegr, Jan Toman

## Abstract

Natural selection is considered to be the main process that drives biological evolution. It requires selected entities to originate dependent upon one another by the means of reproduction or copying, and for the progeny to inherit the qualities of their ancestors. However, natural selection is a manifestation of a more general *persistence principle*, whose temporal consequences we propose to name “stability-based sorting” (SBS). Sorting based on *static stability*, i.e., SBS in its strict sense and usual conception, favours characters that increase the persistence of their holders and act on all material and immaterial entities. Sorted entities could originate independently from each other, are not required to propagate and need not exhibit heredity. Natural selection is a specific form of SBS—sorting based on *dynamic stability*. It requires some form of heredity and is based on competition for the largest difference between the speed of generating its own copies and their expiration. SBS in its strict sense and selection thus have markedly different evolutionary consequences that are stressed in this paper. In contrast to selection, which is opportunistic, SBS is able to accumulate even momentarily detrimental characters that are advantageous for the long-term persistence of sorted entities. However, it lacks the amplification effect based on the preferential propagation of holders of advantageous characters. Thus, it works slower than selection and normally is unable to create complex adaptations. From a long-term perspective, SBS is a decisive force in evolution—especially macroevolution. SBS offers a new explanation for numerous evolutionary phenomena, including broad distribution and persistence of sexuality, altruistic behaviour, horizontal gene transfer, patterns of evolutionary stasis, planetary homeostasis, increasing ecosystem resistance to disturbances, and the universal decline of disparity in the evolution of metazoan lineages. SBS acts on all levels in all biotic and abiotic systems. It could be the only truly universal evolutionary process, and an explanatory framework based on SBS could provide new insight into the evolution of complex abiotic and biotic systems.

## 1 Introduction

### 1.1 Theories on the origin of adaptations

The most important evolutionary discovery of Charles Darwin was probably the identification of natural selection (Darwin, 1860). This process offers the explanation of the origin and accumulation of adaptive, often functionally and structurally complex, characters in organisms. These characters enable organisms to effectively and often sophisticatedly react to the selective pressures of their environment, use its resources, and avoid its detrimental forces. Despite all of this, these adaptations that enable survival and successful reproduction of organisms in complex and changing environments originated through the “primitive” method of trial and error, i.e., without the intervention of any sentient being or existence of a preliminary plan.

Explanations and solutions based on the principle of natural selection were applied in a plethora of other systems in the fields of natural science, technology and even humanities. Over the years, evolutionary biologists discovered that selection has several components and many forms, and that biological evolution is also driven and markedly affected by many other mechanisms, e.g. genetic drift, genetic draft, evolutionary drives, gene flow, and species selection (see e.g. Mayr, 2003). It was also demonstrated that numerous adaptive traits did not originate as biological adaptations but, exaptations, or even spandrels (see e.g. Gould, 2002). Moreover, the complex nature of genetic inheritance, various forms of non-genetic inheritance, and the evolution of multi-level meta-adaptations (such as the ontogeny of metazoans) that affect the evolvability of lineages and canalize their ontogeny and anagenesis returned to the focus of evolutionary and developmental biologists in the last years (see e.g. Laland et al., 2015).

However, natural selection is probably a manifestation of a more general law that affects all material and immaterial entities in the universe, does not require replication and inheritance, and is usually called *survival of the stable*, according to the remark in the first chapter of Dawkins’ book Selfish Gene (Dawkins, 1976, p. 13^1^). At first, it sounds like a tautology: Changeable entities change, whereas stable or rapidly emerging entities accumulate and predominate in the system. Indeed, the claim that the most stable (or persistent) entity lasts the longest time is undoubtedly an axiom (Grand, 2001, p. 34-38; Pross, 2012; Shcherbakov, 2012; Pascal and Pross, 2014, 2015) and this “law” thus seems utterly trivial, at least in a simple model. However, in the real world, coexisting entities interact in a complex manner and the consequent evolution of systems of interacting entities with variable and context–dependent persistence is all but simple (while still characteristic of the perpetual search for states of higher stability) (see e.g. Bardeen, 2009, or Pross, 2003, 2004, 2012; Wagner and Pross, 2011; Pascal and Pross, 2014, 2015, 2016, and references therein). As Shcherbakov (2013) concludes: “This principle – “survival of those who survive” – sounds as a tautology, but it is *the great tautology*: Everything genuinely new emerges through this principle.”

Remarks analogical to Dawkins’ *survival of the stable* were made also by several other researchers (e.g. Lotka, 1922a, b; Simon, 1962; Wimsatt, 1980; Van Valen, 1989; Michod, 2000; Grand, 2001; Maynard Smith and Szathmáry, 2010) whereas possible relations between natural selection and various forms of self-organization were analysed by Weber and Depew (1996). However, to our knowledge, Addy Pross and his colleagues elaborated the idea most profoundly (see e.g. Pross, 2003, 2004, 2012; Wagner and Pross, 2011; Pascal and Pross, 2014, 2015, 2016). The phenomenon itself is very general and probably applies to all fields that concern any form of biological or non-biological evolution. Researchers that touched it from various angles during their investigations called it e.g. *natural selection in the non-living world* (Van Valen, 1989), *survival in the existential game* (Rappaport, 1999; Slobodkin and Rapoport, 1974), *contraction* (Slotine and Lohmiller, 2001), *Persistence Through Time of a lineage* (Bouchard, 2008; Bouchard, 2011), *thermodynamic stability* (Pross, 2003, 2004, 2012; Wagner and Pross, 2011), *the selection of long-lasting structures* (Shcherbakov, 2012), *sorting on the basis of stability* or *sorting for stability* (Flegr, 2010, 2013), *natural selection through survival alone* (Doolittle, 2014), *viability selection* or *selection on persistence* (Bourrat, 2014), *persistence principle* (Pascal and Pross, 2014, 2015, 2016), eventually *ultrastability* (Bardeen and Cerpa, 2015). This loose conceptual embedding is probably related to the fact that only a few theoretical researchers (at least in the field of evolutionary biology) attribute great importance to this phenomenon. For example, Okasha (2006, p. 214), who comments on the topic more thoroughly, calls this phenomenon *weak evolution by natural selection*. According to him, this process cannot generate interesting adaptations and thus he considers it to be (in contrast with *paradigmatic evolution by natural selection*) uninteresting from the evolutionary viewpoint. Godfrey-Smith (2009, pp. 40 and 104), presents a similar opinion. He considers such an extension of the term “natural selection” (i.e., *low-powered Darwinian process*) essentially possible but artificial and basically useless. The opposite opinion has been much rarer. It was explicitly presented, e.g., by Bouchard (2011), Doolittle (2014) or Bourrat (2014). Bourrat (2014) even demonstrated that this process can lead to some class of adaptations in numerical models of evolution. He stated that it could actually stand on the very beginning of biological evolution—original non-replicating entities differing only in their persistence could transform into genuine replicators by the means of this process.

In this paper, we argue that this evolutionary mechanism, which is currently underappreciated and mostly is not taken into account in efforts to explain the origin of characters of living organisms at all, acts upon all biotic and abiotic systems that undergo evolution. In fact, this process may be responsible for a wide range of adaptive traits. In the reaction to its weak conceptual embedding, we propose to call this *survival of the stable* (Dawkins, 1976, p. 13) or, more exactly, temporal manifestation of *persistence principle* (Pascal and Pross, 2014, 2015, 2016), i.e., the general tendency for more stable, persistent and unchangeable entities and characters in the system, unambiguously stability-based sorting (SBS) according to the conception proposed by Vrba and Gould (1986) and Gould (2002, p. 659). This term avoids any connotations that attribute the phenomenon only to material, immaterial, living or non-living entities, its confusion with natural selection, which we consider a specific manifestation of this universal principle (see section 2.1), and its confusion with sorting based on any other kinds of criteria. We will clarify the relationship of SBS and selection more thoroughly in the next section. More particularly, we will show that selection is just one special manifestation of the general process of SBS (a relationship that was implied by numerous evolutionary-biological scholars of the role of persistence in nature mentioned above, e.g. Dawkins, 1976, Okasha, 2006, Godfrey-Smith, 2009, Bouchard, 2011, Doolittle, 2014, or Bourrat, 2014). However, despite being related in their essence, selection, as a special case of SBS, has markedly different evolutionary consequences. Therefore, because the aim of this article is predominantly to demonstrate and stress the different evolutionary consequences of the two processes (deeply understudied SBS in the strict sense and usual conception, and its special case, selection), we will consider SBS and selection as separate phenomena from now on.

## 2 Results and Discussion

### 2.1 The relationship between selection and SBS

All forms of selection, including species selection, require selected entities to originate in reproduction or copying (and thus have an ancestor–descendant relationship) and exhibit at least some degree of inheritance of ancestor qualities (Gould, 2002; Okasha, 2006; Godfrey-Smith, 2009). SBS, on the other hand, does not require any of this. It takes place in all systems with history, i.e., evolution in the broad sense. SBS acts upon all material and immaterial entities regardless of their origin, even entities that originate independently of each other such as snowflakes, cosmic objects during the history of universe, memes, or mutually isolated civilizations. According to the fact that—by definition—unstable and changeable entities expire or change into something else whereas the stable and invariable entities persist, more and more increasingly stable variants of sorted entities accumulate in the system over time, whereas less stable variants gradually vanish. This is true even in the case that less stable entities originate more often in a studied system than their stable variants.

SBS and selection act both in open and growing systems, and in closed systems with a stagnating number of entities. For example, in the course of a snowstorm, the number of competing entities (snowflakes) is not limited and will constantly grow in the snow cover (an open system into which new snowflakes constantly arrive from the system’s surroundings). In such a system, the number of less stable entities will constantly decline, but never reach zero because of the constant share of unstable variants among newly arriving snowflakes^2^. In a closed system, e.g. during the evolution of our universe after the Big Bang with a limited amount of matter available to form space objects, or during memetic evolution of some exclusive religious beliefs that is limited by the number of members of society, more stable entities will gradually replace less stable entities (space objects or memes). The same applies to selection. In an open system, e.g. an exponentially growing unlimited population, the number of individuals better adapted to their environment will gradually grow, but worse adapted individuals will remain in the system too. In a closed system, e.g. in a chemostat or a turbidostat (Flegr, 1997), worse adapted individuals with lower speed or effectiveness of reproduction are quickly displaced by their better adapted counterparts. Thus, in both cases, evolution will proceed faster in closed systems.

In most systems, SBS acquires solely the form of competition among entities for the highest *static stability*, i.e., lowest probability of expiration or transformation of individual entities or their traits into something else. In a particular class of systems—those in which new entities originate from parental entities and inherit their traits—SBS becomes predominantly the competition for the highest *dynamic stability* (Pross, 2003, 2004, 2012; Wagner and Pross, 2011; Pascal and Pross, 2014, 2015, 2016). The competition of stable coexisting entities for the longest static persistence becomes competition for the ability to produce the highest number of their own copies (i.e. the copies of the information how to copy itself), or more precisely, competition for the largest difference between the speed of generation and expiration of these copies. This difference is based both on the longevity of entities (e.g. length of the reproductively active life in organisms), as in the case of static stability, and on the speed of their generation, e.g. reproduction or speciation (*Malthusian kinetics* of Pascal and Pross, 2014, 2015, 2016; see also Pross, 2003, 2004, 2012, and Bourrat’s, 2014, models). Darwin’s natural selection (as well as Dawkins’ interallelic competition, Dawkins, 1986, and Vrba’s and Gould’s species selection, Vrba and Gould, 1986; Gould, 2002) are thus special cases of general SBS. Sorting based on dynamic stability (i.e. selection) and sorting based on static stability differ in the nature of what is sorted—entity itself versus the *information* how to create its copies. From a certain perspective, information emancipates from matter in the case of selection (Shcherbakov, 2012). This makes us to expect both kinds of sorting to take place in evolution of systems of replicating entities with heredity, directly affecting its course and perpetually interacting in their effects.

This is in full agreement with Bourrat’s (2014) arguments that were supported by numerical models of the continuous transformation of populations of entities sorted purely on the basis of static stability to populations of genuine replicators. Similar views were presented even earlier, see e.g. Slobodkin and Rapoport (1974), Rappaport (1999), Bouchard (2008, 2011) or Bardeen (2009). Pross (2003, 2004, 2012), Wagner and Pross (2011) and Pascal and Pross (2014, 2015, 2016 and references therein) studied the role of stability in nature from another angle, differentiating physical forces standing behind stability kinds. Their concept and terminology, however, differ in some important details from the presented one (see Fig. 1 and Appendix).

**Fig. 1.**
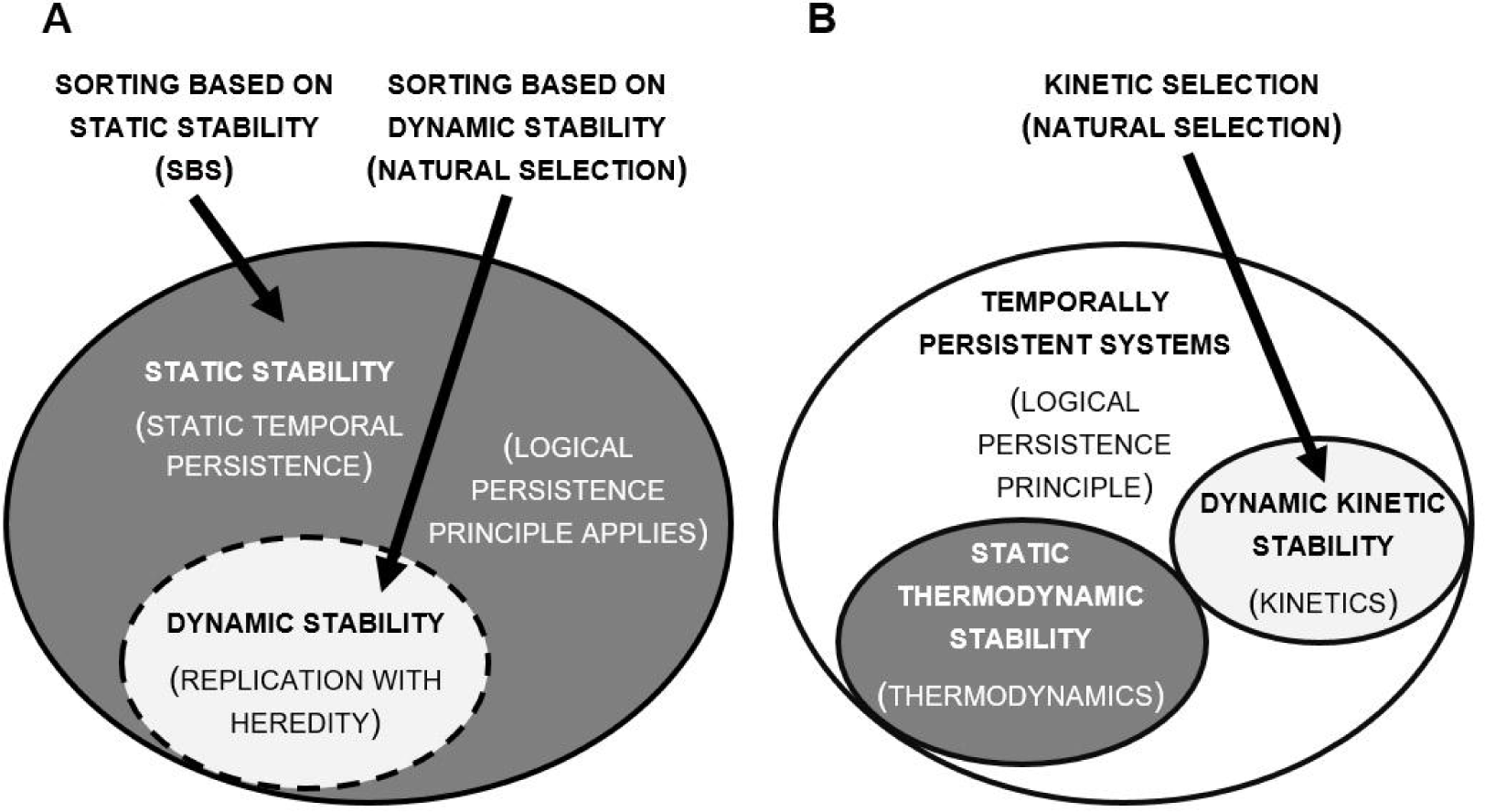
The difference between presented stability concept (A) *and* the stability concept of Press et al. (B). We differentiate two kinds of stability (A). Static stability equates to the entity’s static stability m time, i.e. its persistence until its expiration or ^change^ into something else, regardless of the physical basis of this process Statically mare stable entities and their properties are sorted in time in the process of SBS_ Entities replicating with heredity are sorted, or selected, on the basis of dynamic stability, i.e. largest difference between the speed of generation and expiration of their copies. Putting aside its physical basis and viewed from the evolutionary perspective, however, dynamic stability is only a special case of static stability in systems of entities replicating with heredity in which the statically sorted“thing” became the *information* how to copy itself Pross (2003, 2004, 2012), Wagner and Press (2011) and Pascal and Press (2014, 2015, 2016), on *the* other hand. (B), differentiate static thermodynamic stability and dynamic kinetic stability_ Both of these stability kinds, in the state of high entropy and the exponential multiplication of entities, are governed by the general logical “persistence principle”: systems’ tendency to change from less stable (persistent) to more stable (persistent). Nate that other kinds of stable systems may eventually most and be subject to *the persistence principle_* Dynamic kinetic stability equates dynamic stability in the first concept kinetic selection indeed was proposed to be equal to natural selection (Press, 2004, 2012). It is its relationship to static stability that differs among the two concepts_ Note, (1) that our approach is more general, addresses all material and immaterial entities, and does not address the physical basis of stability, and therefore (2) the difference is mainly conceptual—both approaches need not exclude each other.

In the case that selection, not only SBS in its strict sense, affects the evolution of a certain population; entities that do not invest in their maintenance (and thus have low longevity) but channel the majority of their resources to reproduction may easily prevail. Selection thus represents sorting based on dynamic stability, i.e., a specific form of SBS in the broad sense, whereas SBS in the strict sense and its usual conception represents sorting on the basis of static stability. Therefore, we will respect a traditional terminology, use the term SBS exclusively to refer to sorting on the basis of static stability, and call sorting on the basis of dynamic stability by its standard term—selection (for a more radical approach regarding the classification of selection, see e.g. Pross, 2004, 2012).

It would be erroneous to consider SBS a process from whose direct influence the entities undergoing natural selection completely escaped. As Dawkins (1976, p. 13) stressed, this process is in each sense more general. It acts constantly and simultaneously on all levels. Moreover, the stable accumulates and unstable vanishes regardless of the origin of sorted entities or the nature of the sorting process. Shcherbakov (2012, 2013) goes even further and argues on this basis that the inevitable consequence of every evolution is stasis. Invariance, not variability, is the attractor of evolution. According to this author, any evolutionary changes are only by-products of evolution, e.g. the inability of organisms to completely avoid mutations, or transient consequences of opportunism of selection-based evolution manifested by transient predominance of entities that are less stable in the long-term but have higher dynamic stability—higher fitness—in the short-term. This conclusion might seem quite extravagant taking into account all the variability of life forms on Earth. However, it is the logical consequence of the appreciation of the role of SBS in evolution. It is also worthy to note that Wagner and Pross (2011) and Pross (2012) take the opposite stand, reducing the role of static thermodynamic stability (see Appendix) in the systems of replicating entities only to a general constraint and postulating their general tendency to complexify.

Contrary to both of these approaches, we believe that the role of SBS in the systems of replicating entities with heredity is direct but subtle and selection is rather its tool than by-product, which was suggested only implicitly by Shcherbakov (2012)^3^. In a simple case (stable and homogenous environment), entities in the system would compete only for the highest number and accuracy of copies, i.e., the speed of reproduction associated with its precision, achieved, for example, by reduction of genomic size (which is also the outcome of numerous computer simulations of biological evolution, see e.g. Ray 1993, 1997; Thearling and Ray, 1994, 1996, or Ray and Hart, 1998, as well as experiments, see e.g. Mills et al., 1967). In the real world, the entities are affected by much more heterogeneous conditions of the environment, including other co-evolving entities that undergo selection and mutually interact in a very complex manner. The outcome is constant tension between the pressure to conserve information (i.e., to increase the speed and precision of reproduction) and its evolution (i.e., adaptation to new conditions). Entities that reproduce most rapidly and precisely are not necessarily most successful under these conditions. The increased persistence of individual entities remains the ultimate attractor, yet not by trivial means (static persistence or speed of reproduction), but through more sophisticated adaptations. From our point of view, evolvability is not a mere by-product of evolution. It is an important meta-adaptation that enables an actual increase in the persistence of entities in the process of sorting on the basis of dynamic stability—selection.

Moreover, in the case of terrestrial life, the selected information, which was originally coded directly in the replicating sequence of nucleotides, emancipated to some degree from its material basis. Replicators evolved interactors—bodies—that interpret the information embedded in the sequence of nucleotides in various context-dependent ways. These interactors started new rounds of competition on higher levels, so that the meaning or interpretation of genetic information and the DNA–body complex became the subject of selection rather than the nucleotide sequence itself (Markoš, 2002; Ostdiek, 2011; Shcherbakov, 2012). The consequence is that interacting entities themselves (replicators), as well as the replicated information, change in the course of evolution but still maintain their historical individuality. The outcome of any such competition can be estimated with the help of game theory, particularly the theory of evolutionarily stable strategies (Maynard Smith and Price, 1973; Kolokoltsov and Malafeyev, 2010, p. 65), and if the whole system is complex enough (as e.g. the terrestrial biosphere), it need not immediately follow the path to evolutionary stasis. This, however, does not contradict the SBS-mediated accumulation of stable entities that resist selective pressures and have decreased evolvability; it continuously proceeds on all levels regardless of the effects of selection. The course of evolution on the largest scale can thus be seen as a constant struggle between stability or conservation on one side, and adaptation on the other, which, as will be shown in section 3, can have interesting evolutionary consequences.

### 2.2 Differences between selection and SBS

SBS is much more widespread than natural selection and probably takes place in all evolving systems (i.e., systems with memory/history) with the exception of closed systems with a fixed maximum number of entities in which it proceeded completely, i.e., where only absolutely stable non-expiring entities that are incapable of any change accumulated and remained. In selection, the most successful entities are those that produce the most offspring until their expiration, i.e., death. In SBS, the most successful entities are the most stable ones—those that persist for the longest time without expiring or changing into something else. Selection is much more efficient. Ensuring that offspring inherit the traits of their parents and that the speed of offspring production is based on the number of beneficial traits of the individual, selection gradually accumulates and amplifies beneficial traits, which give individuals a higher dynamic stability—higher fitness. Thus, more (on average) better-adapted individuals and fewer worse-adapted individuals are produced in time. This pattern may be partially masked by the Red Queen effect (Van Valen, 1973). Competitors, predators and parasites evolve counter-adaptations so that, for example, the final speed or effectiveness of reproduction of members of a certain population or species seemingly stagnates until we artificially prevent the counter-evolving species to respond to evolutionary moves of the studied species (see e.g. Becerra et al., 1999). On the other hand, the same share of stable and unstable entities (e.g. snowflakes) originate in the course of SBS regardless of the previous evolution of the system, and especially regardless of the average stability of entities currently constituting the system. This does not fully apply to some memes. For example, new ideas are created with regard to past ones and authors of new ideas preferentially generate such that they have a higher chance of success in long-term competition with existing ones (a process analogical to “copy-the-product”, see Blackmore, 1999, pp. 59-62). However, this is probably specific to entities created by conscious beings that are able to (at least partially) anticipate future development of the system (see e.g. Blackmore, 1999).

In the course of the evolution of a certain genealogical lineage, incomparably more complex adaptations originate by means of the gradual accumulation of mostly small changes (beneficial mutations) in natural selection than by means of much more widespread SBS. It is clear that random changes that increase the stability (persistence) of entities may also accumulate in systems without selection, but this process would be incomparably less effective and slower (see Bourrat, 2014). However, it is possible in principle, as was modelled by Doolittle (2014). In the course of selection in closed systems (which are, in the long term, all systems undergoing biological evolution), every beneficial change spreads to most or even all members of the population. Newly originated beneficial change thus would almost always affect the individuals that already bear the previous one. In SBS, the probability of a simultaneous occurrence of several changes that increase the stability of one newly originated individual is negligible, and the time necessary for the accumulation of a larger number of changes that are beneficial in terms of stability in one individual might be incomparably longer than its estimated lifespan (see Bourrat, 2014). For example, the chamber eye evolved multiple times independently by means of natural selection (Fernald, 2000). It is, however, very unlikely that such a complex organ would evolve solely by means of SBS.

In spite of lower efficiency of SBS, a certain category of adaptations that we see in modern organisms probably originated by means of SBS rather than selection. However, these can only be characters that originated by one or two changes, not a long chain of consequential changes that would continuously elaborate a certain function. An important source of adaptations that increase the stability of sorted entities (e.g. individuals in natural, i.e. intraspecific, selection or evolutionary lineages in species selection) are preadaptations. Such characters evolved by means of selection as adaptations to a certain function, but later turned out to be advantageous in terms of stability and thus spread and prevailed by the means of SBS. SBS works as a sieve that selects characters contributing to the long-term stability of entities that constitute the system and also the system itself (Doolittle, 2014). An example of such a character may be obligate sexuality (Davison, 1998; Flegr, 2008, 2010, 2013; Shcherbakov, 2010, 2012, 2013; Gorelick and Heng, 2011), which originated by natural selection, likely as one of the mechanisms of reparation of mutations, especially structural damage to DNA (Bernstein and Bernstein, 2013; Hörandl, 2013)^4^. Only e*x post* did it turn out that sexuality significantly contributes to the stability of its holders—sexual species—in heterogeneous, changeable and often unpredictable conditions ruling on most of the Earth’s surface. Asexual species are constantly at risk of adapting to temporarily changed conditions, losing their genetic polymorphism and not being capable of re-adaptation to original (or any other) conditions fast enough. This could even lead to their extinction. Sexual species, on the other hand, adapt to transient environmental changes only imperfectly, and constantly maintain high genetic polymorphism (including currently disadvantageous alleles) because of the effects of genetic epistasis and pleiotropy in conjunction with frequency dependent selection. Therefore, they are always able to quickly re-adapt by the changes of allelic frequency (Williams, 1975, pp. 145-146, 149-154, 169; Flegr, 2008, 2010, 2013). From the perspective of individual selection, sexuality is, accompanied by the two-fold cost of meiosis, two-fold cost of sex and other handicaps of its holders (Lehtonen et al., 2012), disadvantageous. From the perspective of species selection—in this case the lower probability of extinction of species in heterogeneous environment—it is highly advantageous. However, species selection is weaker and cannot act against individual one. From the perspective of SBS, it is highly advantageous as well; species and lineages that reverse to asexual reproduction are sorted out, i.e., perish, those that cannot reverse to asexual reproduction for any reason accumulate, and by this mechanism sexuality might spread and prevail.

SBS cannot gradually generate such spectacular adaptations as, e.g., chamber eye, yet it always has the final word in evolution and is even able to completely reverse the course of evolution driven mostly by selection. For example, the human brain and consciousness are undeniably one of the most remarkable characters among terrestrial organisms. However, it is possible that this brain that enabled humans to dominate the Earth and establish a multi-billion population may also be the reason of our early extinction, either by the means of catastrophic warfare, failed physical or biological experiment or “prosaic” severe viral infection that could spread only in a sufficiently dense and interconnected population. From the macroevolutionary point of view, humans may be easily outlived by species in which some ontogenetic constraints in the role of preadaptation prevented the evolution of a sufficiently efficient brain.

Selection is opportunistic. It would beat seemingly “forward planning” SBS in a stable environment (see e.g. Ray, 1993, 1997; Thearling and Ray, 1994, 1996; Ray and Hart, 1998). However, in a changing environment, i.e., under the realistic conditions of Earth’s surface, it is otherwise. Selection does not “plan in advance” and thus is only able to improve the adaptation of organisms on the current conditions regardless of the risk of impairing their future chances of survival, including the risk of extinction of the whole species. Considering the “adaptive landscape” (Wright, 1932), species and populations are able to move only in the upward direction under normal conditions and thus are able to occupy only local, not global, optima. Descending a little and then ascending on another slope for the occupation of a higher peak in the adaptive landscape would not be possible under the normal regime of selection. Mutants that descend have lower fitness and they or their offspring are removed from the population before accumulating other mutations, reaching the “bottom of the valley” and starting to ascend on another slope. On the other hand, SBS does not have such a limitation and is subject to much less opportunism^5^. In the case that a certain adaptation (e.g. a certain pattern of altruistic behaviour) decreases the chance of survival of an individual or slows down its reproduction, yet simultaneously enhances the chances of survival of the population of the individual’s species, those (probably rare) populations and species in which the adaptation prevailed would prosper and survive in the long term.

In most species and within them in most populations, individual selection would act against these tendencies and prefer mutants that lose the individually disadvantageous character. However, populations and species that are preadapted with safeguards against such reverse changes would prevail in the end. Returning to the previous example, such safeguard against the reversal of asexual reproduction may be for example mammalian genomic imprinting that significantly reduces the chance of successful transition to asexual reproduction (Bartolomei and Tilghman, 1997). This and all similar safeguards originated as preadaptations, i.e., adaptations for another purpose, or as spandrels, i.e., characters without any function formed purely as the consequence of topological, physical, biochemical or ontogenetic constraints (see e.g. Gould, 2002). Many species presumably did not have any such safeguards, but we don’t see them today because they lost to their counterparts in the process of SBS. Rare extremes are usually more important than average values both in intraspecific and interspecific evolution (see e.g. Dobzhansky, 1964, or Williams, 1975). Winner usually “takes all”. The fact that the vast majority of populations do not have safeguards and are dominated by selfish individuals means nothing if a safeguard is present in at least some populations. It would be the populations that bear the safeguard that would determine the evolution of a studied species. Similarly, if there happens to be a safeguard against the loss of sexuality or altruistic behaviour in certain species that is absent in the vast majority of others, we will meet only the species with such a safeguard and their descendants in the long term.

## 3 General Discussion

### 3.1 Phenomena in which SBS plays an important role

#### 3.1.1 Microevolutionary phenomena

SBS is much more widespread than selection. However, in the reign of biological evolution, and especially in the processes operating on the human (ecological to microevolutionary) timescale, its significance is obscured by spectacularly manifesting natural selection. SBS is thus encountered especially in phenomena whose origin, establishment or maintenance wasn’t convincingly explained by natural selection yet. Such products of SBS may be, for example, sexuality mentioned in section 2.2 or some types of altruistic behaviour, including restrictions on individual reproduction under the risk of overpopulation that were widely discussed in the past (Dawkins, 1976; Wilson, 1983; Wynne-Edwards, 1986; Leigh, 2010). The usual assumption is that individuals that “voluntarily” reduce the speed of their reproduction would be displaced by selfish mutants (i.e., eliminated by selection). The whole phenomenon is interpreted not as an evolutionary adaptation that increases the long-term success of populations, but as an individual adaptation that enables the individual to save its resources in the situation of high offspring mortality. The proximate reasons for this phenomenon are also being emphasized, e.g. territoriality, social hierarchy, or that individuals in too dense a population disturb each other, reducing the success of each other’s reproduction (Wynne-Edwards, 1986). However, these proximate reasons may act as the safeguards described in section 2.2 that enables certain populations not to be dominated by selfish individuals, which are able to reproduce regardless of the risk of overpopulation. The existence of a safeguard, e.g. the population density-dependent ability “to be disturbed” by nearby individuals, might give the species a chance to overcome the risks of fatal overpopulation and thus give it the decisive advantage in SBS. Species without this or some similar safeguard were more susceptible to extinction and thus we do not meet them today.

Doolittle (2014) suggested that another product of the process that we call SBS may be widespread and often intensive horizontal gene transfer (HGT). According to this author, it may significantly accelerate the adaptations of (especially prokaryotic) organisms to environmental stressors. Such acceleration is probably advantageous in two ways: in terms of individual selection in the short to medium-term and, as will be shown in section 3.1.3, in the long-term because of the gradual stabilisation of environmental conditions (Markoš, 1995; Doolittle, 2014). In a similar way to sexual reproduction mentioned in section 2.2, the original purpose of HGT was probably completely different (it probably served for horizontal spread of selfish genetic elements, see e.g. Redfield, 2001). However, species and lineages that evolved safeguards against the loss of ability to undergo HGTs preserved the ability of relatively fast reactions to the changes of conditions. The most profound safeguard against the loss of HGT ability may be the extraordinary conservation of genetic code (Markoš, 1995; Syvanen, 2002; McInerney et al., 2011)—evolutionary lineages that deviated too much and compromised their ability to undergo HGTs were sorted out by lineages that could still enjoy its benefits.

Similarly, SBS can explain the wide distribution of certain strikingly restrictive traits of modern organisms, i.e., safeguards against the loss of a trait that is beneficial in the long-term. Some examples might be e.g. genomic imprinting of mammals mentioned in section 2.2 or a similar phenomenon in gymnosperms, whose embryos require organelles from the paternal gamete for successful development (Neale et al., 1989); or the extraordinary conservation of genetic code that may enable mutual compatibility of organisms in horizontal gene transfers (Markoš, 1995; Syvanen, 2002; McInerney et al., 2011).

#### 3.1.2 Macroevolutionary phenomena

SBS may also explain certain macroevolutionary phenomena. It is probably tightly connected to the phenomenon of evolutionary stasis, or the punctuated pattern of evolution of (especially) sexual organisms (see e.g. Eldredge and Gould, 1972, or Gould, 2002, pp. 745-1024, with particular examples on pp. 822-874). As was already mentioned, sexual reproduction spread and is still maintained by means of SBS—it helps to maintain high genetic polymorphism, prevents opportunistic one-way adaptation accompanied by loss of genetic polymorphism and enables fast and reversible evolutionary reactions to fluctuations of conditions in changeable and heterogeneous environments by means of epistasis and pleiotropy interconnected with frequency-dependent selection (Flegr, 2008, 2010, 2013). Another consequence of SBS in sexual species is the accumulation of functionally interconnected alleles on the level of an individual and a population. Alleles that are tightly and non-trivially interconnected in their effects on a phenotype, especially alleles that are maintained in a polymorphic state by frequency-dependent selection, are extremely hard to fixate or eliminate through any type of selection and thus are more persistent and accumulate in populations (Flegr, 2008, 2010, 2013). Such “microevolutionary freezing” may be beneficial even to individual organisms—for example, it may enhance the robustness of development to internal and external changes (Shcherbakov, 2012). Sexual species thus remain in evolutionary stasis for most of their existence and are able to irreversibly change only under certain conditions, as was suggested by Eldgredge and Gould (1972)^6^. This is in accordance with Sheldon’s (1996) theory *Plus ça change* that highlights the difference between paleobiological evolutionary patterns of species of changeable environments (punctuated evolution) and species of stable environments (gradual evolution). The difference between these “generalists and specialists in geological timescale” may stem from the presence, or absence, of sexual reproduction.

The very prominent and almost universal pattern of macroevolutionary processes is also a non-monotonous change in disparity, i.e., morphological and functional variability (e.g. in the number of body plans), in the course of the evolution of particular evolutionary lineages, or more precisely, particular taxa. Every clade of an evolutionary tree originates in a speciation event and initially contains a single species. Thus, it has minimal diversity (number of species) and minimal disparity at the beginning. The number of species and morphological and functional diversity then grow in the course of the evolution of a lineage, as do the number of different phenotypically distinct clades and number of higher taxa described by paleotaxonomists within the original evolutionary lineage. However, individual sub-clades die off in time and only clades whose species differ in continuously decreasing number of still less essential traits originate within the remaining clades. The number of species of the original taxon, diversity, need not necessarily decrease and may even grow for a considerable time. Its disparity, on the other hand, decreases (Rasnicyn, 2005; Erwin, 2007; Hughes et al., 2013). According to the class of developmental explanations of this phenomenon, the taxon exhibits high evolvability, i.e., “evolutionary plasticity”, at the beginning. Its members can initially change in almost every trait under appropriate selective pressures. In time, more and more traits “macroevolutionarily freeze”, so that modern members of the taxon are not able to evolve such profoundly new adaptations and lifeforms that were evolved by the species in earlier stages of the evolution of the clade (Foote, 1997; Eble, 1998; Erwin, 2007). The taxon thus gradually abandons different parts of morphospace and perhaps only one, often very specialized and phenotypically very uniform, clade survives at the end. For example, only the species-rich but morphologically rather uniform clade of birds (Aves) survived from original highly disparate clade of dinosaurs to the present (Chiappe, 2009). An even more extreme example of gradual loss of disparity, which is in the long-term probably accompanied by the loss of diversity because of decreasing evolvability, may be the so-called “living fossils” (see e.g. Lloyd et al., 2012). The phenomenon of dead clade walking (Jablonski, 2002), i.e., higher susceptibility to extinction in many isolated lineages of higher taxa that survived mass extinction, may also be a manifestation of the same process. It is probable that these lineages are macroevolutionarily frozen and their possible responses to selective pressures of the post-extinction environment are thus very limited.

A spectacular example of macroevolutionary freezing is the evolution of multicellular animals. The common ancestor of all bilaterians lived approximately 700 million years ago, whereas the common ancestor of all metazoans probably did not precede them by more than 100–200 million years (Douzery et al., 2004; Peterson et al., 2008; Erwin et al., 2011). However, metazoans did not exhibit any significant diagnostic characters until Cambrian or at least Ediacaran, and they probably consisted of mm-sized creatures without hard parts that would enable their identification and classification in fossil material. However, something happened in the early Cambrian approximately 540 million years ago, and metazoans started evolving rapidly and differentiating into many morphologically and ecologically distinct forms, future metazoan phyla (Shu, 2008). This initial period was short and lasted tens of millions of years maximally (Erwin et al., 2011). All current animal phyla, and several tens of other phyla that gradually died out in the next millions of years, originated during this time (Gould, 1989). No other animal phylum and, with the exception of certain groups of radically simplified parasitic organisms (Canning et al., 2004; Glenner and Hebsgaard, 2006; Murchison, 2008), no radically new body plans have originated since the end of the Cambrian. The trend of a gradual decrease of disparity in the course of the evolution of a lineage was also documented in many particular taxa of multicellular animals and plants (Erwin et al., 1987; DiMichele and Bateman, 1996; Eble, 1999). Other examples were summarized by Gould (1989) or Erwin (2007), and, according to Hughes et al. (2013), this trend is characteristic for Phanerozoic clades of metazoans in general. Particular macroevolutionary frozen traits are, for example, the patterns of head segmentation characteristic of main groups of arthropods, five-fingered legs of tetrapods, or (with a few exceptions) seven cervical vertebrae of mammals. All these currently frozen traits were, in some cases even considerably, changeable in the early stages of the evolution of respective taxa (Hughes et al., 2013).

The gradual macroevolutionary freezing of individual traits is almost certainly not just taxonomic artefact caused by the subjectivity of our view from the recent perspective and the way paleotaxonomists delimit taxa of higher and lower level (older combinations of characters delimit higher taxa and *vice versa*). Freezing of individual traits in the course of macroevolution is a real phenomenon that is observed even on the intraspecific level. On this level, it was first described by Italian zoologist Daniele Rosa, and is known as Rosa’s rule today (Rosa, 1899). For example, intraspecific variability of particular morphological characters and the number of characters in which this variability is exhibited are much greater in the early branched-off species than in later branched-off species of certain taxon. Particular evidence for this pattern is the gradual decrease of intraspecific variability in trilobites (Trilobita). Webster (2007) documented that the relative proportion of species with at least one intraspecifically polymorphic morphological character decreased from 75% in middle Cambrian to 8% in late Cambrian. After the subsequent rise to 40% in early Ordovician, it just more or less monotonically decreased until middle Devonian. At that time, the intraspecific polymorphism vanished completely, not to show again until the extinction of taxon at the end of Permian. The temporal pattern in proportion of characters coded as intraspecifically polymorphic is even more striking, declining from a median of 2,8% and 3,6% in middle and late Cambrian to a median of 0% in post-Cambrian. The primary reason for the freezing of individual characters in the course of macroevolution is therefore most likely the freezing of these characters within particular species. If species cease to vary in certain trait, there are no diverse variants of this trait among which selection might differentiate. Such species are thus unable to adapt to conditions to which species cleaved early in the evolution of respective taxon were able to adapt easily (Webster, 2007).

Frozen Evolution Theory (do not confuse with Frozen Plasticity Theory which describes the causes of alternation of short “evolutionarily plastic” and long “evolutionarily elastic” phases in species’ lifetimes) assumes that the reason for the macroevolutionary freezing of individual traits and, consequently, taxa (monophyletic sections of the evolutionary tree delimited by a taxonomist on the basis of a unique combination of several diagnostic characters) of sexual organisms is SBS (Flegr, 2008, 2010, 2013). Various characters exhibit various degrees of evolvability, i.e., the ability to change under appropriate selective pressures, given by the way of their genotype–phenotype mapping and frequency-dependent effect on fitness (Flegr, 2008, 2010, 2013). In the initial phase of the evolution of a certain taxon, a large part of the characters of its members are easily changeable, a smaller part harder and only a small fraction, probably those that the members of the taxon inherited from their evolutionary ancestors, not at all or to a very limited extent. Individual characters change in the course of the taxon’s evolution, even in terms of their variability and evolvability. Traits that are able to change easily and distinctly under proper selective pressures appear and disappear, whereas stable traits persist and accumulate in the taxon. More and more traits irreversibly freeze by means of this sorting, both on the intraspecific and interspecific level. Intraspecific variability is decreasing in a growing fraction of traits. The disparity of the whole taxon is decreasing because old evolutionary lineages of the taxon slowly die out and newly originating species can be distinguished from the original species only to a limited degree in a small and constantly decreasing number of traits.

Organisms, or their evolutionary lineages, may theoretically avoid irreversible macroevolutionary freezing through species selection (Stanley, 1979). Competition for the highest rate of speciation and lowest rate of extinction should theoretically ensure that the lineages with the highest (remaining) evolvability prevail in the long-term. However, SBS, whose manifestation is also macroevolutionary freezing, probably cannot be reversed by species selection, i.e., sorting on the basis of dynamic stability at the species level. Irreversible macroevolutionary freezing is a ratchet-like process that continuously accumulates stable characters and traits in all lineages simultaneously. It cannot be ruled out that certain new species may rarely acquire a unique combination of characters that was not sorted on the basis of stability yet, which would probably mostly accompany its transition to a completely different environment or the adoption of a new ecological strategy. A certain seemingly irreversibly frozen character, or combination of characters, may also exceptionally “thaw” in the course of the evolution of a lineage and start to respond to selection again. Both these events might stand on the beginning of the evolution of birds whose common ancestor conjoined several unique adaptations (Brusatte et al., 2014) and uncoupled the development of front and rear legs to a considerable degree (Dececchi and Larsson, 2013). However, a more fundamental thaw, e.g. thawing of whole body plan, is probably extremely rare, and if it happens, it has the character of a radical simplification of current individual development. This can be demonstrated, e.g., on the example of endoparasitic crustaceans from the clade Rhizocephalia (Glenner and Hebsgaard, 2006), seemingly unicellular endoparasitic cnidarians from the clade Myxozoa (Canning et al., 2004) or sexually- or biting-transmitted mammalian cancers (Murchison, 2008). These radically simplified species may become founders of entirely new, initially macroevolutionary very plastic, but gradually irreversibly freezing high-ranking taxon.

#### 3.1.3 Ecological and geophysiological phenomena

SBS acts even on the ecosystem level, and, in the largest spatial and temporal scale, on the level of the whole planet. Communities in a newly establishing ecosystem (e.g. after severe fire, deglaciation, or emersion of a new island) undergo ecological succession. With a certain degree of simplification, ecosystems are heading towards an equilibrium state—climax—in which they can stay, or around which they can oscillate, for a considerable time in the absence of significant changes to environmental conditions (see e.g. Walker and del Moral, 2003)^7^. The development of communities towards the stage of climax is of various lengths and complications and the final climax communities may vary according to the character of disturbances, amount of available resources and energy etc. (in other words, a climax community may be a polar growth of lichens, as well as a tropical rainforest). Ecological succession is a multidimensional process and takes place on many levels. It may even lead to significant changes in abiotic conditions of the environment. However, it always follows the rules of SBS. Individual species are sorted based on their persistence in the context of a dynamically changing community. An important component of this persistence is their current ecological success. In the long-term, however, their contribution to the stability of the ecosystem is much more important (Bardeen, 2009). This contribution need not be active and need not be paid at the expense of individual fitness (such a system could be extremely easily invaded by selfish entities). It is, rather, based on the species’ ecosystem function, its by-products and side effects—safeguards on the ecosystem level. Species that unidirectionally change the environment towards the conditions suboptimal for them disappear, whereas species that are incorporated in various negative feedback loops that maintain conditions favourable for them persist. Thus, an ecosystem at an advanced stage of development is usually able to compensate (at least to some degree) for the effects of biotic and abiotic environments that lead it off current balance. However, if the intensity of these effects exceeds a certain threshold, the ecosystem may, sometimes profoundly, change (e.g. after distortion of the ecosystem by an invasive species, or change in the soil pH caused by certain tree species). Such change leads to further change in the course of ecological succession (Walker and del Moral, 2003). Certain changes may be destructive—exceptional cases even on the global scale—e.g. the origin of oxygenic photosynthesis that completely altered global conditions on Earth. Such events are described by the Medea hypothesis, see Ward (2009). However, Medea-class events are probably very rare and organisms are thus able to adapt to the resulting changes with the help of selection on the evolutionary timescale. On the other hand, if the changes exceed a critical threshold, or if they are too fast (this applies more to the catastrophic events of abiotic character, e.g. the impact of large cosmic bodies), they can lead to the extinction of all (at least surface) life on the planet.

The strong version of the Gaia hypothesis (Lovelock, 1979) was rejected by most evolutionary biologists because of its assumption that planet Earth (with the help of terrestrial organisms) actively maintains conditions suitable for life. According to the hypothesis, this “planetary homeostasis” is ensured by a broad array of negative-feedback cycles of chemical elements and energy and Earth thus shows signs of a superorganism. The main argument against it is that the only known possible natural origin of such a purposeful system involves natural selection (Doolittle, 1981; Dawkins, 1982; Gould, 1988). However, the group selection on behalf of a whole biosphere postulated by Lovelock would collapse under the pressure of individual selection favouring selfish individuals. The same is true for species selection. The only other alternative, selection on even higher level—the level of whole planets or biospheres—is impossible for one non-reproducing and non-competing individual (the Earth).

Nevertheless, such a long-term stable system integrated by negative feedback loops might develop through sorting of individual geological, atmospheric and biological entities and processes on the basis of stability, i.e., their contribution to the long-term stability of the terrestrial environment. This contribution is possible to estimate with the help of game theory, or more specifically, the theory of evolutionarily stable strategies (Maynard Smith and Price, 1973; Bardeen, 2009; Kolokoltsov and Malafeyev, 2010, p. 65). Entities and processes that did not contribute to the stability of the system or directly led it out of balance acted only temporarily, whereas the ones that supported the long-term maintenance of stability in the context of other forces accumulated. The main difference from ecological succession mentioned earlier in this section, besides the role of biogeochemical cycles that manifest themselves only on higher spatial and temporal scales, is that it operates on evolutionary, not ecological, timescales and new biological entities enter the system through speciations, not colonizations. In a similar way to ecological succession, entities and processes acting against the establishment of homeostasis might (even substantially) change conditions in the system. Nevertheless, the general SBS-mediated tendency of the system to develop towards higher stability via the accumulation of contextually stable elements affects it all the time, on all levels. The later the system is observed, the more long-term stability supporting entities and processes it accumulates and thus remains in stable states for longer periods (Doolittle, 2014). This agrees with the observed decrease in extinction and speciation rates (Raup and Sepkoski, 1982; Gilinsky and Bambach, 1987; Gilinsky, 1994; Benton, 1995; Alroy, 2008) and accumulation of long-lived genera in the terrestrial biosphere during the Phanerozoic (Rohde and Muller, 2005). Decreasing sensitivity of the ecosystem to the effects of newly arriving species was also observed in certain computer simulations, see e.g. Post and Pimm (1983). Another consequence of SBS is that it is more probable that any such system (Earth, certain exoplanet etc.) will be met in a long-term stable state than in an ephemeral unstable one.

SBS acts on any space body, even lifeless ones, and always leads to the most stable states under current circumstances. The equilibrium among geological, geochemical and atmospheric processes need not be static even on lifeless bodies; it could be dynamic, as was observed, e.g., on Venus or Titan, and continuously evolve in relation to changes of outer and inner conditions. However, only in the case when biological entities with a significant effect on the conditions of the environment take part in this sorting, the whole system is heading towards the long-term stable and negative-feedback-regulated conditions favourable for this specific class of entities. The establishment of biogeochemical cycles (planetary homeostasis) is probably further facilitated by the multilevel character of the sorting of biological entities based on their contribution to long-term stability—they are sorted on all levels simultaneously including the global level. SBS is thus able to explain the accumulation of biological entities and processes that maintain conditions suitable for their own survival with the help of negative-feedback processes without greater difficulties. As in the preceding examples, we should not be surprised that, *ex post*, the whole system looks strikingly non-evolutionary, almost like it was planned. This is the common feature of systems evolved by SBS.

Doolittle (2014) and Bardeen (2009) reached similar conclusions regarding the possibilities of establishing Gaian planetary homeostasis; they also postulated the evolution of a system (Earth) towards more stable states through the accumulation of contextually stable elements. Both these researchers supported their arguments by computer simulations: selection of non-replicating non-competing entities in the first case and Gaian “daisyworlds” in the second. Doolittle (2014) got especially close to our conception of SBS. According to this author, classical adaptations do not originate in this process. It can, however, sort adaptations that originated by means of natural selection. These adaptations thus serve as mutations of a higher level. Bardeen (2009) elaborates the basic idea even further and proves that persistence, i.e., long-term stability, is *de facto* the true fitness. Similar reasoning also lies behind proposals to define fitness as the rate of actual or potential persistence of biological entities (in Bouchard’s words “differential survival through a time of a lineage”) in the context of a system (Bouchard, 2008, 2011). However, this is (at least to a high degree) a direct implication of an even older theory of evolutionarily stable strategies. According to this theory, organisms compete for the highest persistence, or the continuing in an “existential game” (Slobodkin and Rapoport, 1974).

#### 3.1.4 Cultural and other phenomena

SBS-based explanations may be naturally applied even in many non-biological fields that deal with evolving systems. The principle of SBS was described and used as an explanation for numerous phenomena e.g. in the fields of artificial intelligence (Slotine, 1994; Runarsson and Jonsson, 1999), cybernetics (Slotine and Lohmiller, 2001; Slotine, 2003), and even cosmology (Safuta, 2011). Its role is probably even more significant in cultural evolution. SBS is able, e.g., to explain the continuous freezing of social institutions and slowing down of social development: It is possible to change almost everything immediately after the establishment of a society, or a revolution that broke down current organization. However, self-maintaining institutions and forces, whose changes gradually slow down and eventually stop, accumulate in time by means of SBS. Numerous authors have highlighted this aspect of cultural evolution. For example, Kováč (2015, p. 26), stressed the evolution of laws, morals, culture and political arrangements towards greater stability. Charles Sanders Pierce named this aspect of cultural evolution “the origin of habit” and “sedimentation” (see e.g. Eco, 2000). Rappaport interprets evolution as constant struggle to maintain stability that is manifested in cultural evolution by the origin, formalisation and petrification of rituals under whose paradigm the society develops (Rappaport, 1999, pp. 416, 425-437). According to Rappaport, the “aim” of all entities is to persist in the existential game as long as possible. This existential game follows the rules of evolutionarily stable strategies, whereas entities that are most stable in the context of their environment and other interacting entities persist for the longest time (Slobodkin and Rapoport, 1974; Rappaport, 1999, pp. 6-7, 408-410, 420, 422-424). However, in a similar way to biological evolution, cultural evolution also need not unidirectionally lead to absolute stability.

Cultural evolution usually has a punctuated character: the alternation of short periods of dynamic changes with long periods of stasis. Systems theory calls this pattern an alternation of “ultrastability” and “breaks” that occur after the deviation of an ultrastable system beyond the limits of its adaptability, which leads to its rearrangement, whether the systems are biological, economical or technological (Bardeen and Cerpa, 2015). This aspect of cultural evolution was highlighted from another angle by Lotman (2009, pp. 7-18, 114-132), who distinguished the periods of cultural “stasis” and “explosion”. Bardeen and Cerpa (2015) presented many particular examples from cultural, or technological, evolution. Numerous particular examples of the punctuated character of cultural evolution were also presented by Gould (2002, pp. 952-972). Markoš (2014) explicitly pointed out the analogy of this pattern of cultural evolution and biological punctuationalism, particularly the processes described by the Frozen Plasticity Theory. In another article (Markoš, 2015), this author connects the ideas of Pierce, Lotman, Rappaport and Flegr and interprets them as various views of the general property of all semiotic systems (historical systems with evolution, ancestor–descendant relationships, memory and experiences): Original chaos “charged” with possibilities follows one specific trajectory, which is plastically changeable at the beginning, but gradually freezes and passes into the state of stasis characteristic of reversible “elastic” reactions to internal and external influences. According to Markoš (2015), the evolution of all semiotic systems ends either by their expiration, or return to the original state of chaos. The biosemiotician Ostdiek (2011) analogically connects the “solidification” of the meaning of a particular symbol and the transition of a system to a state of evolutionary stasis characteristic of elastic reactions. This author even explicitly emphasizes the Frozen Plasticity Theory and argues for the homology of processes causing microevolutionary freezing and solidification of a symbol (particularly its usage by a bigger population and in a higher number of connotations and interactions with other signs and symbols) or the restoration of its original plasticity (only if the symbol loses most of its original meaning). SBS thus takes place even in cultural evolution, although, because of its specifics, SBS sometimes proceeds there in a slightly different manner than in biological evolution.

### 3.2 Historical dimension

The relatively late discovery of the principle of natural selection is considered one of the greatest enigmas of science. This principle is simple and evident from the modern point of view, yet it was not discovered until the latter half of 19^th^ century, i.e., later than the vast majority of comparably complex and many even more complex processes in other fields (Komárek, 2003, pp. 37-44). One reason for this lateness may be cognitive bias. The human brain is specialized in solving problems of interpersonal relations, and every problem that is not easily translated into such a relation or does not have evident analogies with these relations is disproportionately harder to solve (Cosmides, 1989; Gigerenzer and Hug, 1992). For example, it was demonstrated that only a small fraction of unaware respondents solves the *Wason selection task* (Wason, 1966, 1968) easily and correctly: “You are shown 4 cards labelled A, D, 3 and 8 on the visible faces. You know that each card has a letter on one side, and a number on the other. Which card(s) must be turned over to test whether following statement applies to these 4 cards: *If* a card shows A letter, then there is an odd number on the other side?” On the other hand, if we translate the same task into the question on interpersonal relationships: “There are 4 persons in the bar: one elderly and one young, in which we can’t recognize the nature of their drinks, and two persons of uncertain age, one of which drinks an alcoholic beverage and the second soda. Which of these persons must be controlled by a policeman to test whether the bar serves alcohol to minors?”, it is solved easily and correctly by nearly everyone.

The concept of *sociomorphic modelling* (Komárek, 2009) shows that Darwin’s model of natural selection, which explains the evolution of organisms as the consequence of competition of individuals for the highest fitness, could not have been generally thought of and formulated until 19^th^ century England, in which analogous competition among individual economical subjects led to striking and immensely fast development in industry and society. The process of industrial development based on the prosperity of successful and demise of unsuccessful companies was easily thought of, which greatly facilitated insight into an analogical process among living organisms. It is no coincidence that a more or less identical model of evolution was independently formulated by Matthew (1831), Darwin (1859; Darwin and Wallace, 1858) and Wallace (Darwin and Wallace, 1858) within a few years. It is true that ideas preceding the exact formulation of the theory of natural selection could be traced several decades back (see e.g. Rádl, 2015). However, a similar insight would be much more difficult just 100–200 years earlier—back then, there was almost no substantial industrial development and companies; rather, craftsmen workshops were associated with guilds that guaranteed stable prices and quality of their products, and offered practically the same spectrum of products as they did for centuries (Ogilvie, 2004).

On the other hand, the very same rapid development of the material world that has surrounded us until now might have precluded the identification of another universal process that drives biological evolution—SBS—until recently. It is telling that this process was known already in ancient Greece and some historical models of biological evolution were based exclusively on it. For example, Empedocles formulated a model of the origin of living organisms through random combinations of individual limbs (i.e., organs) (Campbell, 2000). Most organisms that arose this way were not successful or even viable because their randomly combined limbs did not fit together very well. However, some of these organisms proved to be well organized, were successful and persistent, and prevailed. Thus, we cannot exclude the possibility that we will not be able to fully recognize and appreciate the true value of the most universal process that drives the evolution of practically all living and non-living systems until the rapid development of our material world slows down or ceases completely.

### 3.3 Conclusion

Natural selection is neither the only, nor the most general process that drives biological evolution. It is a manifestation of a more general but underestimated *persistence principle* (Pascal and Pross, 2014, 2015, 2016), for whose temporal—and hence evolutionary—consequences we have proposed the name “stability-based sorting”. We believe that this neutral term may enable the unification of different approaches to the study of SBS-related phenomena and facilitate the interconnection of different narrowly focused field-specific studies on this topic with related general theoretical-biological concepts.

Our broad concept of stability that consists of (1) static stability and SBS in its strict sense and usual conception, i.e. the accumulation of temporally persistent unchanging entities and characters, and (2) sorting based on dynamic stability, i.e. selection, being a special case of this phenomenon in systems of entities replicating with heredity (see Fig. 1) has broader scope than any other attempt to study these phenomena in the field of evolutionary biology or related disciplines. Therefore, despite our primary goal was to show the paramount importance of the effects of SBS on various levels of diverse evolutionary systems—a fact that has been practically neglected among evolutionary biologists—our conception may also serve as a new standpoint and universal platform for students of various kinds of evolving systems.

All complex novelties in biological evolution originate from the joint influence of two kinds of SBS in the broad sense, the force that drive the system towards dynamic stability and the force that drive the system towards static stability. The same applies to all natural and artificial systems whose entities multiply by reproduction or copying and exhibit at least some degree of inheritance—e.g., cultural evolution or even simulated systems with those properties. Indeed, there are clear analogies between the SBS-related phenomena observed in various kinds of evolving systems, for example, the punctuated character of their evolution or increasing resistance to change (see e.g. Post and Pimm, 1983; Ostdiek, 2011; Markoš, 2014, 2015). Explanatory framework based on SBS thus could provide new insight into the evolution of any complex system.

In future, simulations that recognize the difference between *static* and *dynamic* nature of the sorting the evolving systems undergo and discriminate the role of these two kinds of sorting under various parameters may significantly contribute to the understanding of the general rules of evolution of any systems, and, consequentially, our theoretical understanding of some of the most profound phenomena of existence—e.g., the nature of life.

## 4 Appendix

### 4.1 The relationship between the presented concept and the conception of Pross et al.

Pross (2003, 2004, 2012), Wagner and Pross (2011) and Pascal and Pross (2014, 2015, 2016 and references therein) studied the role of stability in nature thoroughly, differentiating static thermodynamic stability that affects non-living entities and dynamic kinetic stability that is based on replicative chemistry and characteristic of living entities. The identification of the exact physical basis of the stability kinds is out of scope of this article. However, the equation of static stability to thermodynamic stability, i.e. the state of highest entropy (Pross, 2003, 2004, 2012; Wagner and Pross, 2011; Pascal and Pross, 2014, 2015, 2016), is an evident one. Pross and his colleagues stress that this kind of stability is fundamentally different to dynamic kinetic stability based on replicative chemistry and Malthusian kinetics, whereas the two stability kinds are unified under the umbrella of purely logical *persistence principle*: The general tendency of systems to change from less stable (persistent) to more stable (persistent) forms (Pascal and Pross, 2014, 2015, 2016).

Our conception that integrates all evolutionary systems regardless their physical basis is slightly different (see Fig. 1). In our concept, thermodynamic stability is just one option how to ensure static stability, although it could be speculated whether all other options (regarding e.g. immaterial entities such as memes, or even dynamically stable entities) could be ultimately converted or do naturally converge onto this one. Dynamic stability in our conception is not defined by the physical properties of particular system (i.e. replicative chemistry) either. Although the degree of dynamic stability must depend on the Malthusian kinetics of the dynamically stable entities (it would probably be better to say context dependent evolutionary stability in the sense of evolutionary stable strategies of Maynard Smith and Price, 1973) as in the Pross’ concept, we stress especially the second, somehow “static”, aspect of this sorting—heredity. Dynamic stability in our concept can be explicated as a special case of static stability in which the stable sorted “thing” changed from the entity itself to the heritable information necessary for its copying or reproduction. Therefore, static stability in our conception is more general and *de facto* corresponds to Pross’ general persistence in time or *persistence principle* (see Fig. 1).

## 5 Acknowledgment

We thank Charlie Lotterman for the final revisions of our text. Funding: This work was supported by the Grant Agency of the Charles University in Prague (project no: 578416); and the Charles University Research Centre (UNCE 204004). The funding sources had no role in study design, the collection, analysis and interpretation of data, the writing of the report and in the decision to submit the article for publication.

## Footnotes

1 „Darwin’s ‘survival of the fittest’ is really a special case of a more general law of survival of the stable ( …) The earliest form of natural selection was simply a selection of stable forms and a rejection of unstable ones. There is no mystery about this. It had to happen by definition.”

2 Dynamics of such a system were modelled, e.g., by Doolittle (2014).

3 “Evolution is resistance to entropy, the adaptation to environment being only one of the means of this resistance.”

4 The so called “reparation theories” are only one of many concepts proposed for the origin of sexual reproduction. See e.g. Birdsell and Wills (2003) for other proposed theories of the origin of sexual reproduction. However, the vast majority of them assumes that original purpose of sexual reproduction and the reasons of its subsequent spread and long-term maintenance differ.

5 A certain degree of opportunism can manifest only in SBS ongoing in a closed system. Stable entities that are resistant to current effects of environment, or effects that do not actually affect the system but happen relatively often, could prevail there. In closed systems, this precludes the occurrence of entities that would be more resistant to another, possibly much stronger, effect of environment that happens much less often. On the other hand, SBS ongoing in open systems is not opportunistic at all. Ultimately stable entities always prevail there in the long term.

6 Several alternative hypotheses for the conditions under which species in the state of evolutionary stasis may start to irreversibly respond on selective pressures were suggested already by Eldredge and Gould (1972). However, the transition between the “plastic” and “elastic” phase of the species’ evolution is probably most thoroughly described by Frozen Plasticity Theory, see e.g. Flegr (1998, 2008, 2010). All types of punctuationalistic theories of evolution and proposed conditions for the above-mentioned transition were comprehensively summarized by Flegr (2013).

7 Taking into account the plethora of factors of biotic and abiotic environments that affect terrestrial organisms, it is better to consider the concept of climax as depicted here a simplification; a mobile attractor at best, towards which all ecosystems are usually heading but almost never reach. This, however, does not contradict the general tendency of ecosystems to evolve towards a stable climax stage, i.e., the accumulation of species that maintain stable conditions for their survival in the context of other biotic and abiotic factors.

## References

Alroy, J., 2008. Dynamics of origination and extinction in the marine fossil record. Proceedings of the National Academy of Sciences of the United States of America 105, 11536-11542, doi: 10.1073/pnas.0802597105.

Bardeen, M., 2009. Lessons from Daisyworld. Survival of the stable., Centre for Computational Neuroscience and Robotics, Vol. Ph. D. University of Sussex, Brighton, UK, pp. 93.

Bardeen, M., Cerpa, N., 2015. Editorial: Technological Evolution in Society - The Evolution of Mobile Devices. Journal of Theoretical and Applied Electronic Commerce Research 10, 1-7.

Bartolomei, M., Tilghman, S., 1997. Genomic imprinting in mammals. Annual Review of Genetics 31,493-525.

Becerra, M., Brichette, I., Garcia, C., 1999. Short-term evolution of competition between genetically homogeneous and heterogeneous populations of *Drosophila melanogaster*. Evolutionary Ecology Research 1, 567-579.

Benton, M., 1995. diversification and extinction in the history of life. Science 268, 52-58, doi:10.1126/science.7701342.

Bernstein, H., Bernstein, C., 2013. Evolutionary origin and adaptive function of meiosis. In: Bernstein, H., Bernstein, C., Eds.), Meiosis, Vol. 1. InTech, Available from: http://www.intechopen.com/books/meiosis/evolutionary-origin-and-adaptive-function-ofmeiosis.

Birdsell, J., Wills, C., 2003. The evolutionary origin and maintenance of sexual recombination: A review of contemporary models. Evolutionary Biology, Vol 33 33, 27-138.

Blackmore, S., 1999. The Meme Machine. Oxford University Press, New York.

Bouchard, F., 2008. Causal Processes, Fitness, and the Differential Persistence of Lineages. Philosophy of Science 75, 560-570.

Bouchard, F., 2011. Darwinism without populations: a more inclusive understanding of the “Survival of the Fittest”. Studies in History and Philosophy of Science Part C: Studies in History and Philosophy of Biological and Biomedical Sciences 42, 106-114.

Bourrat, P., 2014. From survivors to replicators: evolution by natural selection revisited. Biology & Philosophy 29, 517-538, doi:10.1007/s10539-013-9383-1.

Brusatte, S., Lloyd, G., Wang, S., Norell, M., 2014. Gradual Assembly of Avian Body Plan Culminated in Rapid Rates of Evolution across the Dinosaur-Bird Transition. Current Biology 24, 2386-2392, doi:10.1016/j.cub.2014.08.034.

Campbell, G., 2000. Zoogony and Evolution in Plato’s Timaeus: The Prescoratics, Lucretius and Darwin. In: Wright, M., (Ed.), Reason and Necessity: Essays on Plato’s Timaeus. Duckworth and the Classical Press of Wales, London, pp. 145-180.

Canning, E., Okamura, B., Baker, J., Muller, R., Rollinson, D., 2004. Biodiversity and evolution of the myxozoa. Advances in Parasitology, Vol 56 56, 43-131, doi:10.1016/S0065-308X(03)56002-X.

Chiappe, L., 2009. Downsized Dinosaurs: The Evolutionary Transition to Modern Birds. Evolution: Education and Outreach 2, 248-256.

Cosmides, L., 1989. The logic of social exchange: Has natural selection shaped how humans reason? Studies with the Wason selection task. Cognition 31, 187-276.

Darwin, C., 1859. On the origin of species by means of natural selection or the preservation of favoured races in the struggle for life. John Murray, London.

Darwin, C., Wallace, A., 1858. On the Tendency of Species to form Varieties; and on the Perpetuation of Varieties and Species by Natural Means of Selection. Journal of the proceedings of the Linnean Society of London. Zoology. 3, 45-62.

Davison, J., 1998. Evolution as a self-limiting process. Rivista Di Biologia-Biology Forum 91, 199-220.

Dawkins, R., 1976. Selfish gene. Oxford University Press, Oxford.

Dawkins, R., 1982. The Extended Phenotype: The Long Reach of the Gene. Oxford University Press, Oxford, UK; New York, USA.

Dececchi, T., Larsson, H., 2013. Body and limb size dissociation at the origin of birds: uncoupling allometric constraints across a macroevolutionary transition. Evolution 67, 2741-2752, doi:10.1111/evo.12150.

DiMichele, W., Bateman, R., 1996. Plant paleoecology and evolutionary inference: Two examples from the Paleozoic. Review of Palaeobotany and Palynology 90, 223-247, doi: 10.1016/0034-6667(95)00085-2.

Dobzhansky, T., 1964. How do the genetic loads affect the fitness of their carriers in *Drosophila* populations? American Naturalist 98, 151-166.

Doolittle, W., 1981. Is nature really motherly? CoEvolution Quarterly 29, 59-63.

Doolittle, W., 2014. Natural selection through survival alone, and the possibility of Gaia. Biology & Philosophy 29, 415-423, doi:10.1007/s10539-013-9384-0.

Douzery, E., Snell, E., Bapteste, E., Delsuc, F., Philippe, H., 2004. The timing of eukaryotic evolution: does a relaxed molecular clock reconcile proteins and fossils? Proceedings of the National Academy of Sciences of the United States of America 101, 15386-15391, doi:10.1073/pnas.0403984101.

Eble, G., 1998. The role of development in evolutionary radiations. In: McKinney, M., Drake, J., Eds.), Biodiversity dynamics: turnover of populations, taxa, and communities. Columbia University Press, New York, pp. 132-161.

Eble, G., 1999. Originations: Land and sea compared. Geobios 32, 223-234, doi:10.1016/S0016-6995(99)80036-9.

Eco, U., 2000. Kant and the Platypus: Essays on Language and Cognition. Houghton Mifflin Harcourt, USA.

Eldredge, N., Gould, S., 1972. Punctuated equilibria: an alternative to phyletic gradualism. In: Schopf, T. J. M., (Ed.), Models in Paleobiology. Freeman, Cooper and Co., San Francisco, pp. 82-115.

Erwin, D., 2007. Disparity: Morphological pattern and developmental context. Palaeontology 50, 57-73, doi: 10.1111/j.1475-4983.2006.00614.x.

Erwin, D., Valentine, J., Sepkoski, J., 1987. A comparative study of diversification events: the early Paleozoic versus the Mesozoic. Evolution 41, 1177-1186, doi:10.2307/2409086.

Erwin, D., Laflamme, M., Tweedt, S., Sperling, E., Pisani, D., Peterson, K., 2011. The Cambrian Conundrum: Early Divergence and Later Ecological Success in the Early History of Animals. Science 334, 1091-1097, doi:10.1126/science.1206375.

Fernald, R., 2000. Evolution of eyes. Current Opinion in Neurobiology 10, 444-450, doi:10.1016/S0959-4388(00)00114-8.

Flegr, J., 1997. Two distinct types of natural selection in turbidostat-like and chemostat-like ecosystems. Journal of Theoretical Biology 188, 121-126, doi:10.1006/jtbi.1997.0458.

Flegr, J., 1998. On the “origin” of natural selection by means of speciation. Rivista Di Biologia-Biology Forum 91, 291-304.

Flegr, J., 2008. Frozen evolution: Or, that’s not the way it is, mr. Darwin - Farewell to selfish gene. Createspace Independent Pub., USA.

Flegr, J., 2010. Elastic, not plastic species: frozen plasticity theory and the origin of adaptive evolution in sexually reproducing organisms. Biology Direct 5, -, doi:ARTN 2 DOI 10.1186/1745-6150-5-2.

Flegr, J., 2013. Microevolutionary, macroevolutionary, ecological and taxonomical implications of punctuational theories of adaptive evolution. Biology Direct 8, doi:10.1186/1745-6150-8-1.

Foote, M., 1997. The evolution of morphological diversity. Annual Review of Ecology and Systematics 28, 129-152, doi: 10.1146/annurev.ecolsys.28.1.129.

Gigerenzer, G., Hug, K., 1992. Domain-specific reasoning: Social contracts, cheating, and perspective change. Cognition 43, 127-171, doi: 10.1016/0010-0277(92)90060-U.

Gilinsky, N., 1994. Volatility and the Phanerozoic decline of background extinction intensity. Paleobiology 20, 445-458.

Gilinsky, N., Bambach, R., 1987. Asymmetrical patterns of origination and extinction in higher taxa. Paleobiology 13, 427-445.

Glenner, H., Hebsgaard, M., 2006. Phylogeny and evolution of life history strategies of the Parasitic Barnacles (Crustacea, Cirripedia, Rhizocephala). Molecular Phylogenetics and Evolution 41, 528-538, doi:10.1016/j.ympev.2006.06.004.

Godfrey-Smith, B., 2009. Darwinian populations and natural selection. Oxford University Press, USA.

Gorelick, R., Heng, H., 2011. Sex reduces genetic variation: a multidisciplinary review. Evolution 65, 1088-1098, doi:10.1111/j.1558-5646.2010.01173.x.

Gould, S., 1988. Kropotkin was no crackpot. Natural History 97, 12-21.

Gould, S., 1989. Wonderful Life: The Burgess Shale and the Nature of History. W. W. Norton & Company, New York, London.

Gould, S., 2002. The Structure Of Evolutionary Theory. The Belknap Press of Harvard University Press, Cambridge Massachusetts, London England

Grand, S., 2001. Creation: Life and how to make it. Harvard University Press, Cambridge, USA.

Hughes, M., Gerber, S., Wills, M., 2013. Clades reach highest morphological disparity early in their evolution. Proceedings of the National Academy of Sciences of the United States of America 110, 13875-13879, doi:10.1073/pnas.1302642110.

Hörandl, E., 2013. Meiosis and the paradox of sex in nature. In: Bernstein, H., Bernstein, C., Eds.), Meiosis, Vol. 1. InTech, Available from: <u>http://www.intechopen.com/books/meiosis/evolutionary-origin-and-adaptive-function-ofmeiosis.</u>

Jablonski, D., 2002. Survival without recovery after mass extinctions. Proceedings of the National Academy of Sciences of the United States of America 99, 8139-8144, doi: 10.1073/pnas.102163299.

Kolokoltsov, V., Malafeyev, O., 2010. Understanding Game Theory: Introduction to the Analysis of Many Agent Systems with Competition and Cooperation. World Scientific Publishing Co. Pte. Ltd., New Jersey, London, Singapore, Beijing, Shanghai, Hong Kong, Taipei, Chennai.

Komárek, S., 2003. Mimicry, aposematism and related phenomena: Mimetism in nature and the history of its study. LINCOM EUROPA, Muenchen.

Komárek, S., 2009. Nature and culture: The world of phenomena and the world of interpretation. LINCOM Europa, München.

Kováč, L., 2015. Closing Human Evolution: Life in the Ultimate Age. Springer, Cham, Heidelberg, New York, Dordrecht, London.

Laland, K., Uller, T., Fellman, M., Sterelny, K., Muller, G., Moczek, A., Jablonka, E., Odling-Smee, J., 2015. The extended evolutionary synthesis: its structure, assumptions and predictions. Proceedings of the Royal Society B-Biological Sciences 282, doi: 10.1098/rspb.2015.1019.

Lehtonen, J., Jennions, M., Kokko, H., 2012. The many costs of sex. Trends in Ecology & Evolution 27, 172-178, doi:10.1016/j.tree.2011.09.016.

Leigh, E., 2010. The group selection controversy. Journal of Evolutionary Biology 23, 6-19, doi: 10.1111/j.1420-9101.2009.01876.x.

Lloyd, G., Wang, S., Brusatte, S., 2012. Identifying heterogeneity in rates of morphological evolution: discrete character change in the evolution of lungfish (Sarcopterygii; Dipnoi). Evolution 66, 330-348, doi: 10.1111/j.1558-5646.2011.01460.x.

Lotka, A., 1922a. Contribution to the energetics of evolution. Proceedings of the National Academy of Sciences of the United States of America 8, 147–151, doi:-.

Lotka, A., 1922b. Natural selection as a physical principle. Proceedings of the National Academy of Sciences of the United States of America 8, 151–154, doi:-.

Lotman, J., 2009. Culture and Explosion. Walter de Gruyter GmbH & Co., Berlin.

Lovelock, J., 1979. Gaia: A New Look at Life on Earth. Oxford University Press, Oxford, UK.

Markoš, A., 1995. The ontogeny of Gaia: the role of microorganisms in planetary information network. Journal of Theoretical Biology 176, 175-180, doi:10.1006/jtbi.1995.0186.

Markoš, A., 2002. Readers of the Book of Life: Contextualizing Developmental Evolutionary Biology. Oxford University Press.

Markoš, A., 2014. Biosphere as semiosphere: Variations on Lotman. Sign System Studies 42, 487-498.

Markoš, A., 2015. The Birth and Life of Species–Cultures. Biosemiotics, 1-12.

Matthew, P., 1831. On naval timber and arboriculture: with critical notes on authors who have recently treated the subject of planting. Adam Black; Longman, Rees, Orme, Brown, and Green, Edinburgh, London.

Maynard Smith, J., Price, G., 1973. The logic of animal conflict. Nature 263, 15-18.

Maynard Smith, J., Szathmáry, E., 2010. The major transitions in evolution. Oxford University Press Inc., New York.

Mayr, E., 2003. The Growth of Biological Thought: Diversity, Evolution, and Inheritance. The Belknap Press of Harvard University Press, Cambridge, Massacusetts; London, UK.

McInerney, J., Pisani, D., Bapteste, E., O'Connell, M., 2011. The public goods hypothesis for the evolution of life on Earth. Biology Direct 6, doi:10.1186/1745-6150-6-41.

Michod, R., 2000. Darwinian Dynamics: Evolutionary Transitions in Fitness and Individuality. Princeton University Press, Princeton New Jersey, Chichester West Sussex.

Mills, D., Peterson, R., Spiegelman, S., 1967. An extracellular Darwinian experiment with a self-duplicating nucleic acid molecule. Proceedings of the National Academy of Sciences 58, 217-224.

Murchison, E., 2008. Clonally transmissible cancers in dogs and Tasmanian devils. Oncogene 27, 19-30.

Neale, D., Marshall, K., Sederoff, R., 1989. Chloroplast and mitochondrial DNA are paternally inherited in S*equoia sempervirens* D. Don Endl. Proceedings of the National Academy of Sciences of the United States of America 86, 9347-9349.

Ogilvie, S., 2004. Guilds, efficiency, and social capital: evidence from German proto industry. The Economic History Review 57, 286-333.

Okasha, S., 2006. Evolution and the levels of selection. Oxford University Press, USA.

Ostdiek, G., 2011. Cast in Plastic: Semiotic Plasticity and the Pragmatic Reading of Darwin. Biosemiotics 4, 69-82, doi: 10.1007/s12304-010-9108-7.

Pascal, R., Pross, A., 2014. The nature and mathematical basis for material stability in the chemical and biological worlds. Journal of Systems Chemistry 5, -, doi:10.1186/1759-2208-5-3.

Pascal, R., Pross, A., 2015. Stability and its manifestation in the chemical and biological worlds. Chemical Communications 51, 16160-16165, doi:10.1039/c5cc06260h.

Pascal, R., Pross, A., 2016. The logic of life. Origins of Life and Evolution of Biospheres 46, 507-513, doi:10.1007/s11084-016-9494-1.

Peterson, K., Cotton, J., Gehling, J., Pisani, D., 2008. The Ediacaran emergence of bilaterians:congruence between the genetic and the geological fossil records. Philosophical Transactions of the Royal Society B-Biological Sciences 363, 1435-1443, doi: 10.1098/rstb.2007.2233.

Post, W., Pimm, S., 1983. Community assembly and food web stability. Mathematical Biosciences 64, 169-192, doi:10.1016/0025-5564(83)90002-0.

Pross, A., 2003. The driving force for life’s emergence: kinetic and thermodynamic considerations. Journal of theoretical biology 220, 396-406.

Pross, A., 2004. Extending the concept of kinetic stability: toward a paradigm for life. Journal of physical organic chemistry 17, 312-316.

Pross, A., 2012. What is life? How chemistry becomes biology. Oxford University Press, Oxford, UK.

Rappaport, R., 1999. Ritual and Religion in the Making of Humanity. Cambridge University Press, New York, USA, pp. 535.

Rasnicyn, A., 2005. Collected works in evolutionary biology (Izbrannye trudy po evolucionnoj biologii). Tovarisevstvo naucnych izdanii KMK, Moscow.

Raup, D., Sepkoski, J., 1982. Mass extinctions in the marine fossil record. Science 215, 1501-1503, doi: 10.1126/science.215.4539.1501.

Ray, T., 1993. An Evolutionary Approach to Synthetic Biology: Zen and the Art of Creating Life. Artificial Life 1, 179-209.

Ray, T., 1997. Evolving complexity. Artificial Life and Robotics 1, 21-26.

Ray, T., Hart, J., 1998. Evolution of differentiated multi-threaded digital organisms. In: Adami, C., et al., Eds.), Artificial Life VI: Proceedings of the Sixth International Conference on Artificial Life. MIT Press, Canbridge Massachusetts, London England, pp. 295-306.

Redfield, R., 2001. Do bacteria have sex? Nature Reviews Genetics 2, 634-639.

Rohde, R., Muller, R., 2005. Cycles in fossil diversity. Nature 434, 208-210, doi:10.1038/nature03339.

Rosa, D., 1899. La Riduzione progressiva della variabilità e i suoi rapporti coll'estinzione e coll'origine delle specie. Clausen, Torino.

Runarsson, T., Jonsson, M., 1999. Genetic production systems for intelligent problem solving. Journal of Intelligent Manufacturing 10, 181-186, doi:10.1023/A:1008928804949.

Rádl, E., 2015. The history of biological theories. BiblioLife, USA.

Safuta, J., 2011. Spacetime deformations evolution concept. arXiv.

Shcherbakov, V., 2010. Biological species is the only possible form of existence for higher organisms: the evolutionary meaning of sexual reproduction. Biology Direct 5, doi:10.1186/1745-6150-5-14.

Shcherbakov, V., 2012. Stasis is an Inevitable Consequence of Every Successful Evolution. Biosemiotics 5, 227-245, doi:10.1007/s12304-011-9122-4.

Shcherbakov, V., 2013. Biological Species as a Form of Existence, the Higher Form. In: Pavlinov, I., (Ed.), The Species Problem - Ongoing Issues. InTech, Rijeka, Croatia, pp. 65-91.

Sheldon, P., 1996. Plus ça change–a model for stasis and evolution in different environments. Palaeogeography Palaeoclimatology Palaeoecology 127, 209-227.

Shu, D., 2008. Cambrian explosion: Birth of tree of animals. Gondwana Research 14, 219-240, doi:10.1016/j.gr.2007.08.004.

Simon, H., 1962. The architecture of complexity. Proceedings of the American Philosophical Society 106, 467-482.

Slobodkin, L., Rapoport, A., 1974. An Optimal Strategy of Evolution. The Quarterly Review of Biology 49, 181-200.

Slotine, J., 1994. Stability in adaptation and learning. In: Cliff, D., (Ed.), From animals to animats 3. MIT Press, Brighton, England, pp. 30-34.

Slotine, J., 2003. Modular stability tools for distributed computation and control. International Journal of Adaptive Control and Signal Processing 17, 397-416, doi:10.1002/acs.754.

Slotine, J., Lohmiller, W., 2001. Modularity, evolution, and the binding problem: a view from stability theory. Neural Networks 14, 137-145, doi:10.1016/S0893-6080(00)00089-7.

Stanley, S., 1979. Macroevolution, Pattern and Process. W.H. Freeman and Company, San Francisco.

Syvanen, M., 2002. Recent emergence of the modern genetic code: a proposal. Trends in Genetics 18, 245-248, doi:10.1016/S0168-9525(02)02647-1.

Thearling, K., Ray, T., 1994. Evolving Multi-cellular Artificial Life. In: Brooks, R., Maes, P., Eds.), Artificial Life IV: Proceedings of the Fourth International Workshop on the Synthesis and Simulation of Living Systems. MIT Press, Cambridge Massachusetts, London England, pp. 283-288.

Thearling, K., Ray, T., 1996. Evolving parallel computation. Complex Systems 10, 229-237.

Van Valen, L., 1973. A new evolutionary law. Evolutionary Theory 1, 1-30.

Van Valen, L., 1989. Three paradigms of evolution. Evolutionary Theory 9, 1-17.

Vrba, E., Gould, S., 1986. The hierarchical expansion of sorting and selection: sorting and selection cannot be equated. Paleobiology 12, 217-228.

Wagner, N., Pross, A., 2011. The nature of stability in replicating systems. Entropy 13, 518-527.

Walker, L., del Moral, R., 2003. Primary Succession and Ecosystem Rehabilitation. Cambridge University Press, Cambridge, UK.

Ward, P., 2009. The Medea Hypothesis: Is Life on Earth Ultimately Self-Destructive? Princeton University Press, Princeton, USA; Oxford, UK.

Wason, P., 1966. Reasoning. In: Foss, B., (Ed.), New Horizons in Psychology, Vol. 1. Penguin Books, Harmondsworth, UK, pp. 135-151.

Wason, P., 1968. Reasoning about a rule. Quarterly Journal of Experimental Psychology 20, 273-281, doi: 10.1080/14640746808400161.

Weber, B., Depew, D., 1996. Natural Selection and Self-Organization: Dynamical Models as Clues to a New Evolutionary Synthesis. Biology and Philosophy 11, 33-65.

Webster, M., 2007. A Cambrian peak in morphological variation within trilobite species. Science 317, 499-502, doi: 10.1126/science.1142964.

Williams, G. C., 1975. Sex and evolution. Princeton University Press, Princeton, NJ.

Wilson, D., 1983. The group selection controversy: history and current status. Annual Review of Ecology and Systematics 14, 159-187, doi:10.1146/annurev.es.14.110183.001111.

Wimsatt, W., 1980. The units of selection and the structure of the multi-level genome. In: Asquithand, P., Giere, R., Eds.), PSA: Proceedings of the Biennial Meeting of the Philosophy of Science Association, Vol. 2. Philosophy of Science Association, East Lansing, MI, pp. 122-183.

Wright, S., 1932. The roles of mutation, inbreeding, crossbreeding, and selection in evolution. In: Jones, D., (Ed.), Proceedings of the Sixth International Congress on Genetics. Brooklyn botanic garden, New York, pp. 356-366.

Wynne-Edwards, V., 1986. Evolution Through Group Selection. Blackwell Scientific Publications, Oxford.

